# Centimeter-scale perfusable cultured meat with densely packed, highly aligned muscle fibers via hollow fiber bioreactor

**DOI:** 10.1101/2023.09.04.555230

**Authors:** Minghao Nie, Ai Shima, Shoji Takeuchi

## Abstract

The development of in-vitro biofabrication methods for producing cultured meat based on animal cells has been advancing, but replicating the texture of traditional meat in centimeter-scale has been a challenge. To address this, a method using a hollow fiber bioreactor (HFB) has been developed. The HFB contains semipermeable hollow fibers that act as artificial circulatory systems to deliver nutrients and oxygen uniformly to the tissue, along with microfabricated anchors for inducing cell alignment. With active perfusion, the biofabricated centimeter-scale chick muscle tissue shows elevated levels of marker protein expression and sarcomere formation across the whole tissue, along with improved texture and flavor. In the future, further scaling up of this approach using industrial robots has the potential to transform not only the cultured meat industry but also the tissue engineering fields aiming for the formation of large-scale artificial organs.

## Introduction

In light of increasing concerns regarding animal welfare and environmental impact associated with meat, a major source of animal protein, the development of in-vitro biofabrication methods for producing cultured meat has been advancing^1–3^. To be widely accepted, cultured meat needs to replicate both the nutritional composition and the textural characteristics of traditional meat^4,5^. Previous methods for producing aligned muscle tissues, which are crucial for replicating the texture of traditional meat, have relied on populating myoblasts within a hydrogel and fixing the hydrogel with anchoring structures or central pillars to induce the cell alignments^2,6–11^. This method has been successful in producing millimeter-scale contractile muscle tissue that can be used for a variety of applications, including biohybrid actuators^11,12^, disease models^13,14^, implantable grafts^15^, and cultured meat^2,16–18^. However, scaling up the biofabrication of muscles remains a major challenge since the previous methods lack proper schemes to deliver nutrients and oxygen uniformly across larger tissues. In densely cellularized tissues, the diffusion of nutritional molecules can be delayed, leading to necrosis if the tissue is too thick without an integrated circulatory system^19^. Consequently, the thickness of tissues without an integrated circulatory system has generally been limited to less than a millimeter, making it challenging to produce centimeter-scale or larger muscle tissues.

In this work, we develop a method for producing skeletal muscle tissues using a hollow fiber bioreactor (HFB). The HFB contains semipermeable hollow fibers that act as artificial circulatory systems to deliver nutrients and oxygen uniformly to the tissue, along with microfabricated anchors for inducing cell alignment (**Fig. 1**). Using the HFB, we first investigate the application of positive internal pressure within the hollow fibers for promoting the formation of densely packed myotubes. Next, the HFB consisting of an array of 50 uniformly distributed hollow fibers is used to construct centimeter-scale chicken muscle tissues, where the benefits of active perfusion are checked morphologically, functionally, and nutritionally for cultured meat applications.

**Figure 1.**
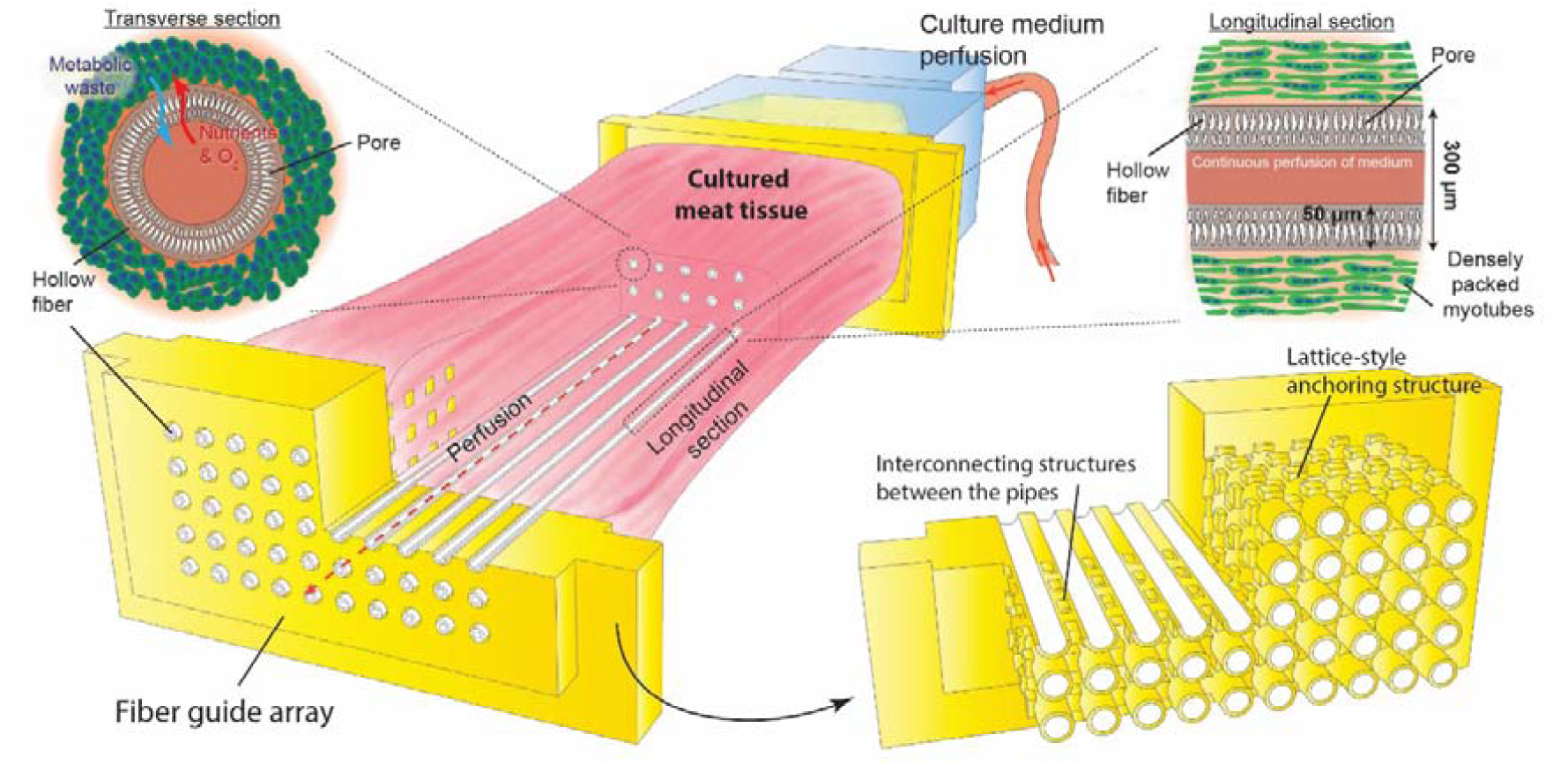
Biofabrication of cultured meat tissue using the hollow fiber bioreactor (HFB). The bioreactor consists of semipermeable hollow fibers which act as artificial circulatory systems to uniformly deliver nutrients and oxygen to the tissue. A key component of the HFB is the microfabricated fiber guide arrays, which are used to define the spatial arrangement of the hollow fibers and to anchor the meat tissue on its two ends. With active perfusion of culture medium, nutrients and oxygen can be delivered to the cells surrounding the hollow fiber while the metabolism waste can be disposed of effectively, therefore achieving the uniform formation of densely packed and alignment myotubes throughout the whole meat tissue.

## Results

### The HFB for the biofabrication of muscle tissues: design and implementation

Our design for a hollow fiber bioreactor (HFB) includes an array of hollow fibers held in place by microfabricated anchors with lattice-style geometry (**Fig. 2**). The fiber guide arrays, fabricated with a high-precision stereolithography machine, feature an array of fiber-threading holes with a fixed pitch distance (i.e. the distance between the centers of two adjacent fibers) of 0.9 mm to position each PolyEtherSulfone hollow fiber (ID: 200∼250 µm, OD: 300∼350 µm). The fibers are further guided by interconnected pipes with a wall thickness of 100 µm to form a lattice-style geometry. During tissue formation, the hydrogel precursors can fill in the vacant spaces between the lattice frames, tethering the tissue to the lattice frames to achieve the anchoring effect. The end of the pipe structures of the fiber guide arrays are designed to be complementary to increase efficiency for fiber threading. Complementary pipe ends of the fiber guide arrays are shown in **Fig. 2e-f**.

**Figure 2.**
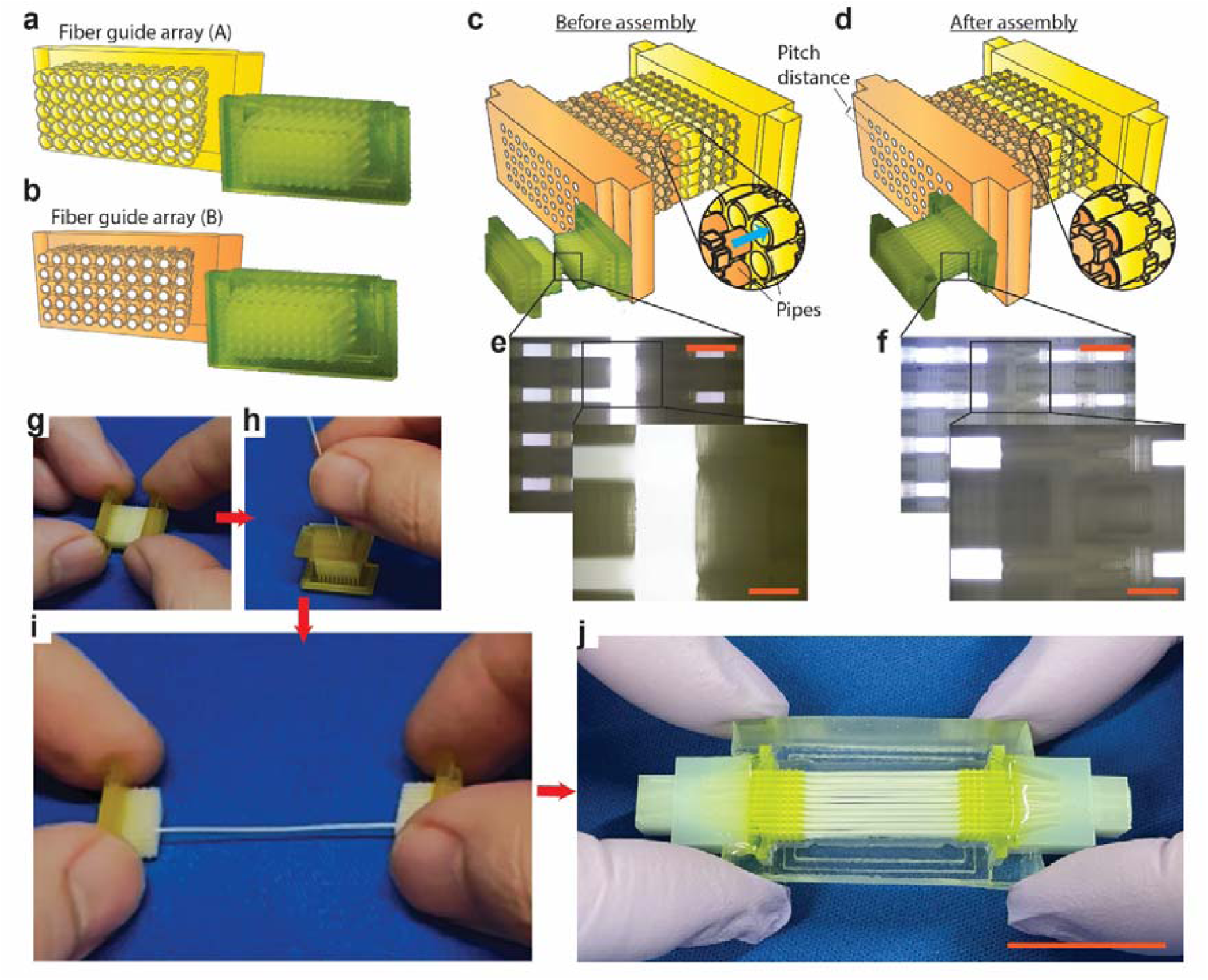
Design and fabrication of the HFB using high precision stereolithography and hollow fibers. **a-b**, Design of the fiber guide arrays (A) and (B), which are featured with arrays of fiber-threading holes with fixed pitch distances and complementary pipe ends that can be pre-assembled to facilitate the threading of the hollow fibers. Photo images of the fabricated fiber guide arrays are as shown in the right. **c-d**, The fiber guide arrays before and after assembly with photo images as shown in the left bottom. **e-f**, Microscopic images featuring the pipe ends of the fiber guide arrays before and after the assembly. **g-i**, Step-by-step procedures for the assembly of the HFB starting with the assembly of the fiber guide arrays, then the threading the hollow fibers, and finally the separation of the fiber guide arrays. **j**, The HFB after the assembly of hollow fibers and the application of gluing. Scale bars, **e-f**:1 mm; inset: 400 µm; **j**: 2 cm.

To assemble the HFB (**Mov. S1**), the fiber guide arrays are assembled (**Fig. 2g**), and the hollow fibers are inserted **(Fig. 2h**). The fiber guide arrays are then separated (**Fig. 2i**), and after gluing and cutting, the fabricated HFB is complete (with 50 hollow fibers, before attaching the perfusable caps) (**Fig. 2j**). Detailed steps for the assembly of the HFBs with various numbers of hollow fibers can be found in Methods.

### Perfusion condition optimization using a single-fiber HFB and C2C12 cells

Experiments were conducted using a simplified HFB with a single hollow fiber (i.e., the “1-fiber HFB”, refer to detailed designs in Methods) to determine optimal perfusion conditions for tissues made with C2C12 cells (an immortalized mouse myoblast cell line). This approach was taken to avoid potential clogging of fibers in the parallel connected array due to bubbles or debris in the culture media, which could result in uneven pressure distribution. By using a single hollow fiber, reliable perfusion within the fiber was achieved, and clogs or bubbles could be removed effectively during perfusion.

First, to determine the optimal perfusion condition for the tissues, we analyzed cell nuclei density and distribution and tissue morphology. Cell nuclei densities were analyzed for tissues cultured for 4 days in growth medium with perfusion flow rates of 15 µL/min, 100 µL/min, and 500 µL/min (the internal pressure difference between the middle point of the hollow fiber and the culture medium outside are calculated to be 0.1 mBar, 0.8 mBar, and 3.7 mBar, respectively; **Table S1**). The tissues were cryosectioned (with the hollow fiber) and stained using DAPI. A representative image of the DAPI-stained transverse tissue section (500 µL/min) is as shown in **Fig. 3a**. High magnification transverse tissue sections images around the hollow fiber for the tissues cultured under control (no perfusion) and perfusion (500 µL/min) conditions are shown in **Fig. 3b** and **Fig. 3c**, respectively. Cell nuclei densities were quantified in various donut-shaped radical regions (defined by an inner radius R_1_ and an outer radius R_2_) (**Fig. 3d**). High cell nuclei densities (>1.0 ×10^3^ cells/mm^2^) were achieved with the 500 µL/min condition, and the nuclei density decreased for areas farther from the center of the hollow fiber. In primitive trials to further increase the internal pressure difference between the middle point of the hollow fiber and the culture medium outside to 50.1 mBar (cultured device: 4-fiber HFB, refer to detailed designs in Methods; **Table S1**), we found the actual withdrawing rate of the culture medium at the device outlet could not keep up with the rate of the syringe, which led to intensive bubble formation inside the device and the syringe.

**Figure 3.**
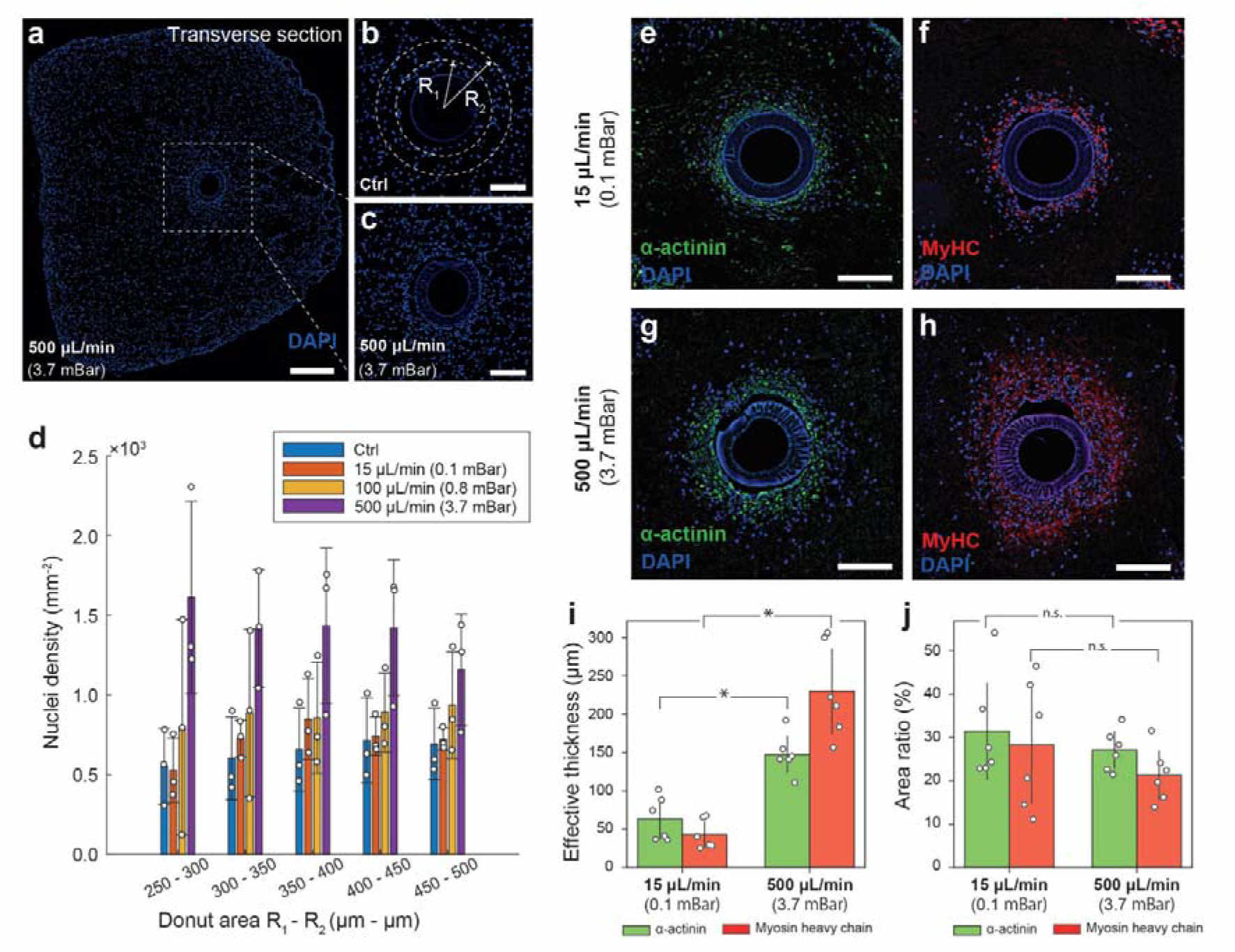
Perfusion condition optimization using a single-fiber HFB. **a**, Transverse section of the tissue with cell nuclei stained with DAPI (blue). **b-c**, High magnification images of the transverse tissue section near the hollow fiber. The tissues were cultured under control condition and perfusion condition (flowrate: 500 µL/min) for **b** and **c**, respectively. **d**, Nuclei density distribution in various radical regions (defined by an inner radius R_1_ and an outer radius R_2_) for the samples cultured under different conditions. **e-h**, Immunostaining of the tissue cross section to indicate the expression of α-actinin and myosin heavy chain (MyHC) for the tissue cultured under perfusion condition with different flowrates (15 and 500 µL/min). **i**, Effective thickness of the donut shape area enclosing all the positively stained cells. Bars represent the mean ± SD. *p < 0.05, Two-tailed, paired, student’s t test (n = 6 stained tissue sections). **j**, The area ratio of the summed area of the positively stained cells against the donut shape area enclosing all the positively stained cells. Bars represent the mean ± SD. Two-tailed, paired, student’s t test (n = 6 stained tissue sections). Scale bars, **a**:500 µm; **b-c**: 200 µm; **e-h**: 200 µm.

Next, tissue morphology was analyzed for tissues cultured for 10 days in growth medium (Day 0-1) and differentiation medium (Day 2-10) with perfusion flow rates of 15 µL/min and 500 µL/min. Myosin heavy chain (MyHC) and α-actinin expression were immunostained, and larger areas of positively stained areas were observed in samples with higher flow rates (500 µL/min) as shown in **Fig. 3e-h**. The adjacent expression areas were merged and enclosed into a donut-shaped area, and the effective thickness of the donuts for both markers (MyHC and α-actinin) are plotted in **Fig. 3i**. The effective thickness of the MyHC and α-actinin expressing areas was more than 2 times larger for the 500 µL/min condition, with an effective thickness 230 ± 61 µm of and 148 ± 26 µm, respectively. The area ratio of the summed expression areas against the merged donut shaped area is 27.1 ± 4.7% and 21.3 ± 6.3% for MyHC and α-actinin, respectively, with no statistically significant differences, indicating a consistent level of myotube density for both the control and the perfusion conditions.

### Biofabrication of centimeter-scale cultured chicken meat

Centimeter-scale (2 cm ×1 cm ×5 mm) chicken meat tissues were fabricated using chick primary myoblasts suspended in Type-I collagen precursors and cultured under bidirectional perfusion (flowrate: 2 mL/min; calculated internal pressure: 38.1 mBar) in a hollow fiber bioreactor consisting of 50 hollow fibers (i.e., the “50-fiber HFB”, refer to detailed designs in Methods) with alternative inject-withdraw cycles (see Methods for details). After 9 days of culture, the cell-laden hydrogel shrank and formed a tissue (**Fig. 4a**) that was cut from the HFB with the hollow fibers embedded (**Fig. S1**), and the hollow fibers can be manually removed subsequently (**Fig. 4b**). Histological analysis was performed on the tissues cultured with and without perfusion (control).

**Figure 4.**
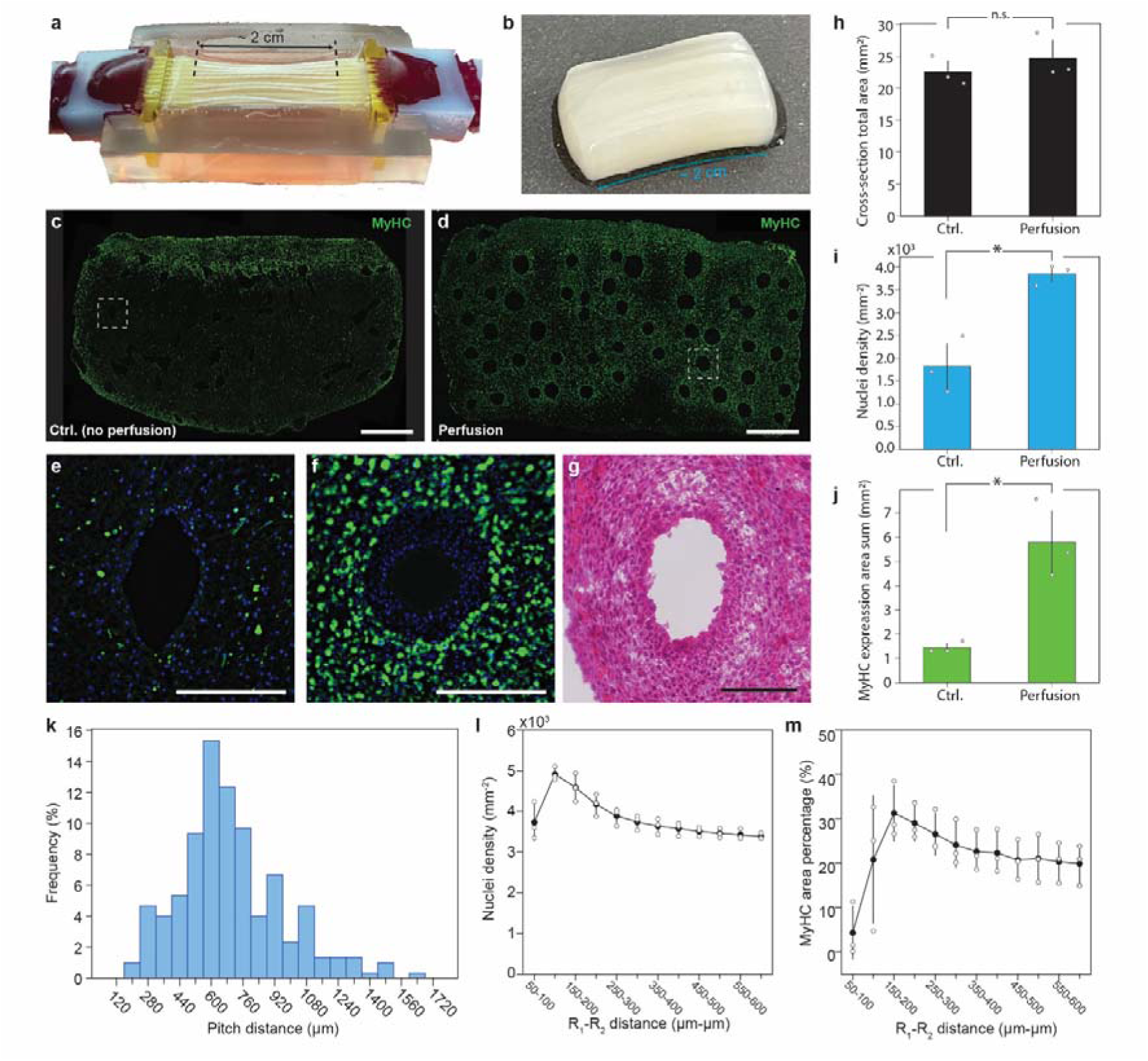
Biofabrication of centimeter-scale cultured chicken meat and transverse section analysis of tissue morphology. **a**, The centimeter-scale chicken meat tissue (2 cm×1 cm×5 mm) fabricated with perfusion culture using the HFB. **b**, The chicken meat tissue after the retrieval of the hollow fibers. **c-d**, Low magnification images of the tissue transverse-sections immunostained using anti-MyHC antibodies for the tissues cultured under control (no perfusion) and (d) perfusion conditions. **e-f**, High magnification images of the immunostained tissue transverse-sections (MyHC: green, DAPI: blue) of the area highlighted using dashed square boxes in **c** and **d**, respectively. **g**, High magnification H&E staining images of the tissue (with perfusion) transverse sections. **h-j**, Quantitative comparison of the tissues cultured under control (no perfusion) and perfusion conditions in terms of transverse section total area, nuclei density, and sum of MyHC expression area, respectively. Bars represent the mean ± SD. *p < 0.05, Two-tailed, paired, student’s t test (N = 3 cultured meat tissues). **k**, Distribution of the actual pitch distances between the adjacent hollow fibers. (mean: 699 µm, standard deviation: 262 µm, CV: 37.5%) **l-m**, Nuclei densities and MyHC area percentages in various radical regions (defined by an inner radius R_1_ and an outer radius R_2_ from the center of the hollow fibers) for the samples cultured under perfusion condition, respectively. Scale bars, **c-d**:1 mm; **e-f**: 250 µm; **g**: 100 µm.

Transverse sections of the tissues were stained using anti-MyHC antibodies and DAPI. The expression of MyHC across the whole cryosections for the control and perfused tissues was shown in **Fig. 4c** and **Fig. 4d**, respectively, while images of the hollows in the dashed regions for the control and perfused tissues were shown in **Fig. 4e** and **Fig. 4f**, respectively. The H&E staining image around a hollow fiber for the tissue cultured with perfusion was shown in **Fig. 4g**. Transverse-section total area, nuclei density, and sum of MyHC positive area for both conditions were shown in **Fig. 4h**, **4i**, and **4j**, respectively, while the distribution of the measured pitch distances between the adjacent hollows for the perfusion cultured tissues was calculated with a mean of 699 µm (CV = 37.5%) and plotted in **Fig. 4k**. Note that PFA fixation and sucrose treatment to acquire tissue cryosections can cause tissue shrinkage (ca. 60%), the actual size of the tissue before fixation should be larger than the numbers indicated in **Fig. 4h**. Nuclei densities and MyHC area percentages for the perfusion cultured tissues were calculated for various radial regions (defined by an inner radius R_1_ and an outer radius R_2_ from the center of the hollow fibers) and shown in **Fig. 4l-m**.

The longitudinal sections of the tissues were stained using anti-α-actinin antibodies and DAPI, which showed the formation of striated patterned sarcomeres of the myotubes, as shown in **Fig. 5a-c**. To quantify myotube alignment, the angles (θ) between the major axes of the fitting eclipses of the hollow fiber and of the myotubes surrounding the hollow fiber were calculated and plotted (**Fig. 5d**). Electrical stimulations were performed to verify the contractile function of the fabricated chicken meat tissue, and the contraction length of the tissues stimulated using alternative ±5.5 V/cm electrical pulses (pulse duration: 20 ms, positive peak-to-peak period: 1 s) was shown in **Fig. 5e**. With active perfusion, the contractile length of the tissues was significantly improved compared to the control tissues (cultured without perfusion) (see SI for **Mov. S2**). Despite that the tissues were fixed with the two anchors and stiffened with the embedded hollow fibers during the electrical stimulation, the contraction at the edges of the perfused tissue was inspectable under a microscope with an amplitude of ca. 5 µm, indicating the successful reconstruction of contractile function of the perfused tissue.

**Figure 5.**
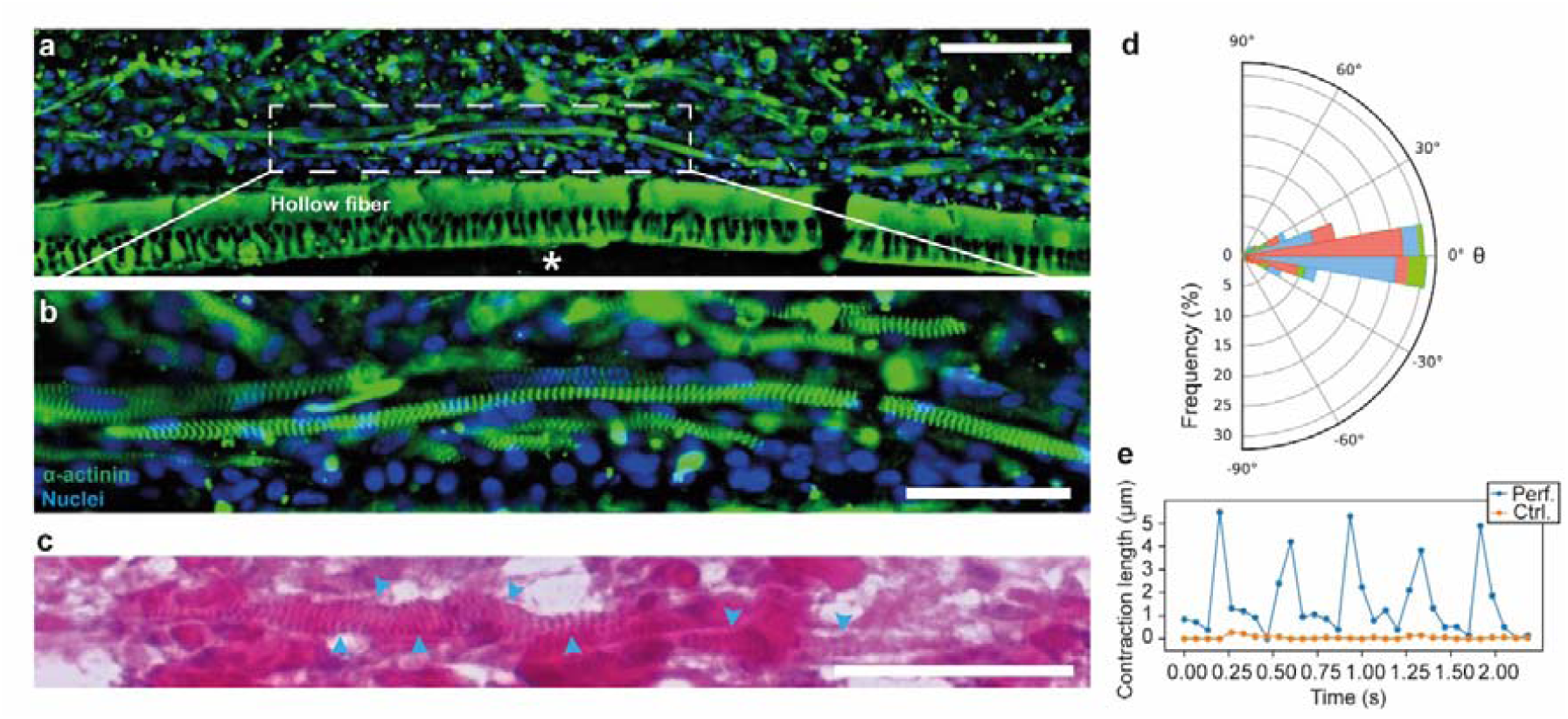
Longitudinal section analysis of tissue morphology and contractile function analysis of the centimeter-scale cultured chicken meat tissues. **a***, Low magnification image of the longitudinal tissue section (*α*-actinin: green, nuclei (DAPI): blue). The asterisk indicates the inner space of the hollow fiber.* **b***, High magnification zoomed image of the longitudinal tissue section of the area highlighted using dashed square boxes in* **a. c***, High magnification image of the longitudinal tissue section stained using H&E with the blue arrows indicating the myotubes with striated patterned sarcomeres.* **d***, Myotube alignment quantified by calculating the angle (*θ*) between the major axes of the fitting eclipses of the hollow fiber and of the myotubes surrounding the hollow fiber. (N = 3 cultured chicken meat tissues, n = 5 hollow fibers with >50 myotubes surrounding each hollow fiber). The histograms for the different tissue samples were colored differently and overlapped in the graph.* **d***, Contraction length of the tissues stimulated using alternative ±9.5V pulse signals (pulse duration: 20 ms, positive peak-to-peak period: 1 s). Scale bars, **a:** 100 µm; **b-c:** 50 µm*.

Texture profile analysis (TPA) and free amino acids (FAAs) analysis were performed for both the perfused and the control tissues. The stress-time curves obtained by running a TPA cycle at room temperature were shown in **Fig. 6a**. The measured stress at 50% strain and tissue half thickness were significantly higher for the perfused tissues compared to the control tissues, as shown in **Fig. 6b**. The FAAs analysis is summarized in the cumulative bar graph shown in **Fig. 6c**, where various types of FAAs were measured and categorized into three major categories with different color tones (red: sweetness, green: umami, blue: bitterness). Compared to the control condition, both the cumulative amount of the FAAs and the fraction of sweetness and umami were higher for the perfused tissue.

**Figure 6.**
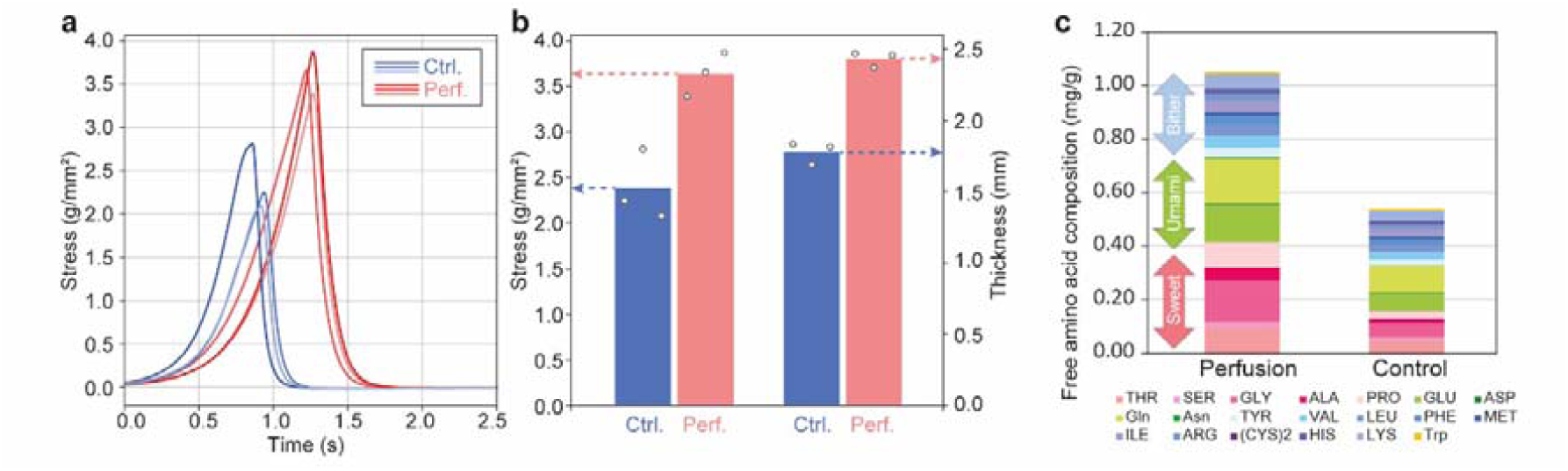
Texture profile analysis (TPA) and free amino acids (FAAs) analysis of the centimeter-scale cultured chicken meat tissues. **a***, Stress-time curves obtained by running a TPA cycle at room temperature (n = 3 cuts of a cultured chicken meat tissue).* **b***, Measured stress (@50% strain) and half thickness of the samples culture under control and perfusion conditions. *p < 0.05, Two-tailed, paired, student’s t test (n = 3 cuts of a cultured chicken meat tissue).* c*, Cumulative bar graph indicating the composition and total amount of free amino acids for both control and perfusion samples. (N = 1 cultured chicken meat tissue)*.

## Discussions

We have proposed a hollow fiber bioreactor (HFB) using high precision stereolithography to achieve the long-term perfusion of large-scale cultured meat. Our experiments confirm that the applied pressures within the hollow fibers not only increase the confluency and promote differentiation activity of myoblasts but also induce muscle contraction capabilities. Furthermore, we successfully demonstrate the biofabrication of centimeter-scale (2 cm ×1 cm ×5 mm) chick skeletal muscle tissues using an array of 50 hollow fibers. With active perfusion, the chick muscle tissue shows significantly higher levels of marker protein expression and sarcomere formation across the whole tissue, along with improved texture and flavor confirmed using TPA and FAAs analysis methods.

The biofabrication of cultured whole-cut meats is a challenging process, as it requires the formation of densely packed, highly aligned muscle fibers across a length scale larger than centimeters. In comparison to the scaffold-based approaches and bioprinting methods which cannot achieve a high cell density uniformly throughout the whole tissue^5,15,20–22^, modular assembly methods have been effective in increasing the cell density in the fabricated tissues^2,11,17,18,23–26^. In detail, first, hydrogel precursors with a high cell concentration (typically > 5.0×10^7^ cells/mL) with a submillimeter cross-sectional size were solidified and cultured to form the microtissue modules. The microtissue modules can avoid central necrosis due to their small cross-sectional sizes^27^. Then, the microtissues were assembled into large avascular meats through tissue fusion^17^ or glueing^18^. However, such modular assembly methods require enormous efforts to develop robotic systems to manipulate and assemble the massive amount of tissue modules at an industrial scale and may also result in weak adhesion between the fused modular tissues, potentially causing deteriorated meat texture and flavor.

To address this challenge, we propose to form a consistent piece of cultured meat tissue with in-situ constructed perfusable channels. By culturing the tissue with active perfusion through vessel-mimicking internal channels, uniform mass transport can be achieved, facilitating the differentiation of myoblasts across the whole tissue. We found that the nuclei and MyHC expression area percentage tend to converge with increasing distance to the center of the hollow fibers (**Fig. 4l-m**), indicating that this approach can lead to the formation of densely packed muscle fibers on a centimeter scale. While the tissues fabricated in this study may not be large enough to meet the market needs for whole-cut meat products, they can already be used as scaled-up tissue modules (in comparison to the millimeter-scale avascular tissue modules^17,18^) to achieve more efficient tissue assembly and reduce the meat-to-glue ratio in the final products. Potentially, the further scaling up of the HFBs used in this work could eliminate the need for tissue assembly altogether. The automation of HFB production and tissue fabrication using industrial robots has the potential to transform the cultured meat industry. Traditionally, the HFB materials for medical applications have been mainly optimized for dialyzers which focus on mimicking the selective permeability of kidneys^28^. For food applications, edible hollow fiber materials, such as cellulose, could also be developed to eliminate the need for removing the fibers after fabrication and create meats with adjustable textures.

Though techniques on embedding perfusable channels in artificial tissues have been previously proposed, these methods require tricky gluing to connect to tubings^29^ and cannot withstand high pressure to mimic in-vivo interstitial flow^30–35^. In contrast, HFBs can be tightly sealed to ensure steady and leak-free perfusion culture^36,37^, and have been used for space-effective mass-culture of cells for artificial organs^38^ and cell therapy^39^ due to their high area-to-volume ratio. However, HFBs have not been successfully applied to form muscle tissues since the current HFBs don’t have anchors to prevent the tissue shrinkage during culture.

Also, filling the hydrogels in between the hollow fibers with uniform spacing is also difficult since the hollow fibers are randomly bundled. In this paper, we have proposed the HFB featured with 3D printed lattice-style anchoring structures and uniformly arranged hollow fibers. First, with the 3D printed lattice-style structures, a unidirectional tension within the cultured tissues has been maintained during culture for the induction of myotube alignment (**Fig. 5**). Also, interestingly, the small angles between the directions of the hollow fibers and the surrounding myotubes hint at the possible effects of the embedded hollow fibers on guiding the cell alignment in 3D skeletal muscle tissues (**Fig. 5d**). Second, with a uniform spatial arrangement of the thin (0.30 -0.35 mm diameter) hollow fibers with a small pitch distance of ca. 699 µm (**Fig. 4k**) enabled by the high-precision stereolithography, a uniform delivery of nutrient and oxygen was achieved to fabricate a consistent and firm tissue. By perfusing culture medium through the closely arranged hollow fibers with an internal pressure of 38.1 mBar, a uniform muscle cell distribution is achieved, which is crucial for the formation of densely packed and highly aligned muscle fibers across a length scale larger than centimeters with improved texture (**Fig. 6b**) and taste characteristics (**Fig. 6c**). Notice that there are also limitations on increasing the internal pressure, as the rate of the culture medium withdrawal at the device outlet may not keep up with the rate of the syringe, leading to intensive bubble formation inside the device and syringe.

The biofabrication of artificial organs that match physiological size has been a longstanding challenge in tissue engineering^40,41^. Our proposed hollow fiber bioreactor (HFB) not only aims to produce large-scale cultured meat but also holds tremendous potential beyond the food industry. Building upon the existing HFBs, we propose further advancements to enhance the capabilities of our HFB system. As one of the possible directions, surface modifications of the hollow fibers could improve cell attachment. To facilitate the formation of heterogeneous tissues, iPS and organoids formation technologies could be adapted to our HFB systems. Since the formation of cell-laden ECMs has been a popular method for tissue formation, many of the current organoid formation protocols can be easily applied to our HFB to enable the perfusion culture of large size organoids^42^. Also, considering the crucial role of mechanical stimuli in tissue development, incorporating stretchable hollow fibers could be a promising avenue for future research.

This breakthrough in creating larger tissues signifies a significant milestone in the field of tissue engineering. The implications extend to various applications, including regenerative and transplant medicine, where the formation of large-scale organs is highly sought after. Additionally, our HFB technology offers opportunities to establish perfusable tissue models for drug discovery and develop thick tissues for biohybrid robots, paving the way for exciting advancements in these interdisciplinary fields. It is also worthy of mentioning that the formation of large-scale artificial organs could not only benefit biomedical research but also robotics fields. For example, the biofabrication of large-scale muscles demonstrated here could lead to the formation of biohybrid actuators with unprecedented power output in comparison to the current millimeter-sized muscle actuators^43,44^.

Currently, our HFB is fabricated by manually inserting each individual fiber, leveraging robotics systems or similar technologies to streamline the process could enable the formation of even larger tissues in the future. However, it is important to acknowledge that as tissue size increases, ensuring sufficient oxygen supply becomes more challenging and will require advanced strategies to effectively dissolve oxygen into the culture mediums.

Nevertheless, we speculate that scaling up could be achieved by exploring innovative approaches such as the development of artificial blood to lift the upper limitation of the conveyable oxygen in the media^45^. In the future, the adaptation and scaling-up of various microfabrication techniques, including photolithography, stereolithography, and multiphoton polymerization^46^, could facilitate the biofabrication of even larger tissues and organs.

Taken together, we have proposed a top-down strategy for the creation of centimeter-scale cultured meat using a hollow fiber bioreactor (HFB). By forming myoblast-laden hydrogels with the embedded, ready-to-perfuse hollow fibers, a consistent piece of muscle tissue can be nurtured uniformly throughout the whole tissue. In comparison to the bottom-up modular assembly methods which use tissue glues to assemble millimeter-scale tissue modules, our top-down strategy has the potential to realize the one-stop biofabrication of whole-cut culture meats with densely packed, highly aligned muscle fibers. With active perfusion, the biofabricated centimeter-scale chick muscle tissue shows significantly higher levels of marker protein expression and sarcomere formation across the whole tissue, along with improved texture and flavor. Further scaling up of this approach using industrial robots has the potential to transform not only the cultured meat industry but also the tissue engineering fields aiming for the formation of large-scale artificial organs.

## Online Methods

### Design and fabrication of the hollow fiber bioreactor (HFB)

#### 1-fiber HFB

As shown in **Fig. S2A-B**, the design of the fiber holder for the 1-fiber HFB adopted an all-in-one style, where the micro anchor arrays (simple cubic lattices consisting of cuboids with 0.4×0.4 mm rectangular cross-section) are designed to grab the tissue during culture and the barbed connector is designed for the easy connection with silicone tubes. **Fig. S2F-G** shows the assembly of the HFB, first, a pair of fiber holders were mounted on a PDMS substrate. The distance between the two fiber holders is designed to be 10 mm. Then, hollow fibers (PolyEtherSulfone hollow fiber, ID: 200-250 µm, OD: 300-350 µm, courtesy of DAICEN MEMBRANE-SYSTEMS Ltd.) were threaded through the holes in the fiber holders. Finally, epoxy adhesives (Araldite RT30, 3M) are added in the reservoirs in the fiber holders to seal the fibers with barbed inlets or silicone tubes and the bioreactors are connected to syringes for perfusion. After the solidification of the adhesives, silicone tubing is connected to the barbed connectors.

#### 4-fiber HFB

The 4-fiber HFB has an array of hollow fibers with a pre-defined pitch distance and a pair of micro anchors with lattice-style geometry to hold the tissue on its two ends. As shown in **Fig. S2C-E**, the design of the fiber holder for the 4-fiber HFB adopted a two-component style, where the micro anchor arrays (simple cubic lattices consisting of cuboids with 0.4×0.4 mm rectangular cross-section) are designed to grab the tissue during culture on one part, and the barbed connector is designed for the easy connection with silicone tubes is designed on another part. After mounting a pair of fiber holders (the part with micro anchor arrays) on a PDMS substrate, hollow fibers (same as the ones used for 1-fiber HFB) were threaded through the 4 pair holes in the fiber holders. Finally, the parts with barbed connectors are assembled and epoxy adhesives (Araldite RT30, 3M) are added in the reservoirs in between the two parts of the fiber holders to seal the fibers with barbed inlets.

#### 50-fiber HFB

Since the increase of the number of hollow fibers requires a more efficient way of fiber threading, the 50-fiber HFB is fabricated by the assembly of various components including one pair of fiber guide arrays ((A) and (B)), two fiber bundle guides, two perfusable caps, one PDMS holder, one removable, as well as the hollow fibers (**Fig. S3**). The key components for enabling the three-dimensional arrangement of the hollow fibers and for anchoring the tissue are the two complementary fiber guide arrays ((A) and (B)), which are fabricated using a high precision stereolithography machine (BMF). As shown in **Fig. S3**, both fiber guide arrays are featured with an array of fiber-threading holes with as-designed pitch distances to define the spatial position of each fiber. Each threaded fiber is further guided by separate pipes that are interconnected to form a lattice-style geometry. With the lattice-style geometry, the hydrogel precursors for forming the tissue can fill into the vacant spaces between the lattice frames and therefore tether the tissue with the lattice frames to achieve the anchoring effect. To efficiently thread the hollow fibers (same as the ones used for 1-fiber and 4-fiber HFBs) through the two fiber guide arrays, the end of the pipe structures is designed to be complementary so that the fiber guide arrays can be assembled with their fiber threading holes pre-aligned. The fiber guide arrays are separated after all the fibers are successfully threaded through and set onto a PDMS holder. Next, the fibers are bundled and further threaded to the fiber bundle guides where the fibers will be further fixed together by the application of epoxy glue (Araldite RT30, 3M). At the open end of the fiber bundle guide, the glued fiber bundles are further cut and trimmed to expose the cross sections of the hollow fibers. The fiber cross-sections in **Fig. S4** are randomly arranged since the fibers are randomly bundled and threaded through the fiber bundle guide. The precise spatial arrangement of the fibers can only be maintained between the two fiber guide arrays. Color differences in the fiber holes are due to the angle of inspection, with some holes appearing dark when the embedded fiber sections align with the inspection angle.

Changing the angle of inspection shows that all holes are not clogged, as the white holes turn dark. Then, the open end of the fiber bundle guide was inserted into a perfusable cap with a barbed connector for the application of silicone tubing, and epoxy glue is applied to seal the cap with the fiber bundle guide. To finish the assembly of the HFB, a removable cap is inserted to cap the bottom of the PDMS holder. After the assembly, the cavity for the application of hydrogel precursors can be defined as surrounded by the two fiber guide arrays, the PDMS holder, and the removable cap. To prevent liquid leakage from the cavity during the crosslinking of the hydrogel, complementary ridges and trenches are designed on both the fiber guide arrays, the PDMS holder, and the removable cap so that their boundaries after assembly can become watertight.

### Cell preparation, tissue construction and perfusion culture

#### C2C12 cells culture and tissue construction

Mouse myoblasts cell line, C2C12 were purchased from American Type Culture Collection (ATCC) and cultured in Dulbecco’s Modified Eagle Medium (DMEM) (high-glucose) (Nacalai Tesque) containing 10% fetal bovine serum (FBS) (Thermo Fisher Scientific) and 0.5% penicillin-streptomycin (Merck) (growth medium; GM) at 37 °C in 5% CO_2_ to obtain the required number of cells. For tissue construction, C2C12 cells at passage 6 were collected and suspended in porcine type I collagen, Cellmatrix Type I-A (Nitta Gelatin) mixed with an equal amount of Matrigel (Corning) at 5 ×10^7^ cells/mL. Cellmatrix Type I-A was neutralized according to the manufacturer protocol. The cell-hydrogel mixture was poured into 1-fiber HFB (ca. 350 µL) and crosslinked in a humidified cell incubator at 37 °C for 30 min (**Fig. S2H**). The tissue was then cultured in GM for 4 days with (15, 100, and 500 µL/min) or without (as a control) perfusion for the growth condition assay, and in GM for 1 day and in differentiation medium (DM) for 9 days with perfusion (15, and 500 µL/min) for the differentiation condition assay. DM was DMEM (low-glucose) (FUJIFILM Wako Pure Chemical) containing 10% horse serum (Thermo Fisher Scientific), 100 ng/mL IGF-I (R&D Systems), 200 µM L-ascorbic acid phosphate magnesium salt n-hydrate (Wako Pure Chemical Industries), and 0.5% penicillin-streptomycin. The medium was exchanged every 2-3 days.

#### Chick primary myoblasts culture and tissue construction

For the chick primary skeletal muscle cell culture, fertilized chicken eggs were purchased from Keikou Sangyo (Ibaraki, Japan) and embryonic development was promoted at 37.6 - 39.5 °C in a poultry incubator (Showa Furanki) for 10 - 12 days. After the chick embryos were taken out from the eggs and washed in phosphate buffered saline (PBS), breast and thigh muscles were dissected. The muscles were minced, washed in PBS, and digested by 0.5% trypsin-EDTA (Merck) at 37 °C for 30 min. The digested tissue was then suspended in chicken culture medium (CCM) and myoblasts were collected through cell strainers after centrifuge. The chick primary myoblasts were plated on tissue culture dishes coated by 1% gelatin (Merck) and cultured in CCM for a day before used for the tissue construction. CCM was DMEM (high-glucose) containing 12% horse serum, 4% chicken serum (Thermo Fisher Scientific) and 0.5% penicillin-streptomycin. This animal experiment was approved by the Office for Life Science Research Ethics and Safety of The University of Tokyo and carried out according to their guidelines. For the chicken muscle tissue construction, the cells were suspended in Cellmatrix Type I-A at 1.5 ×10^8^ cells/mL and 1.5 mL of the cell-hydrogel mixture was poured into the 50-fiber HFB. After being crosslinked for 30-45 min in a humidified cell incubator at 37 °C, the tissue was cultured in CCM for 9 days with perfusion (40 µL/min per hollow fiber) or without perfusion. The medium was exchanged every 2-3 days.

#### Perfusion culture method

After tissue formation, the HFBs are flipped upside down, placed in a culture dish, and connected to a syringe through a via drilled on the lid of the dish. Alternative bidirectional flows are generated by the pull-push motion of the syringe pumps (push/pull volumes: 6/6 mL, flowrate is changed for different culture conditions and can be found in **Table S1**). The medium level of the dish is kept below 1 mm to ensure oxygen saturation level during the perfusion culture. The outlet tube of the HFB is dipped towards the bottom of the culture dish to ensure successful suction of culture medium during the pull phase of the syringe pump. Note that one side of the HFB is connected to an outlet tube, which has a L-shape and is pointed towards the bottom of the culture dish during culture (**Fig. S2I**). The length and the inner diameter of the outlet tube can be tuned according to **Table S1** for various perfusion conditions.

### Cryosectioning and staining of the tissue sections

#### Staining cell nuclei for the growth condition assay

For counting the number of C2C12 cell nuclei under the growth condition, the tissue was taken out from the HFB by cutting both ends with a blade, fixed by 4% paraformaldehyde (PFA), embedded in O.C.T. compound (Sakura Finetek Japan) and frozen in liquid nitrogen for cryosectioning. The sections perpendicular to the hollow fiber (transverse sections) were cut to a thickness of 10 µm by using Cryostar NX50 (Thermo Fisher Scientific), attached to adhesive glass slides (CREST coated slides) (Matsunami Glass), and sealed by VECTASHIELD Mounting Medium with DAPI (Vector Laboratories), staining cell nuclei with DAPI.

#### Immunostaining for myogenic markers

For immunostaining for myogenic markers expressed under the differentiation condition, the tissues were fixed by 4% PFA after ejecting from HFB, the hollow fibers were removed by tweezers if necessary, and incubated in sucrose solutions (10 wt% in PBS(-) for > 2 hrs @RT, 30 wt% in PBS(-) for > 2 hrs @RT) before frozen. The sections parallel to the hollow fiber(s) (longitudinal sections) (thickness: 20 µm) as well as transverse sections (thickness: 10 µm) were prepared. For immunostaining, the sections were incubated in 1% bovine serum albumin in PBS for blocking before being treated with antibodies. Anti-α-actinin antibody (EA-53, Merck) and anti-myosin heavy chain antibody (MF20, Developmental Studies Hybridoma Bank) were used as primary antibodies and Alexa Fluor 488 or 594 goat anti-mouse IgG (Thermo Fisher Scientific) was used as a secondary antibody. The stained sections were sealed by VECTASHIELD Mounting Medium with DAPI.

#### Hematoxylin and eosin (H&E) staining

For H&E staining, both transverse and longitudinal sections were cut to a thickness of 10 µm and attached to MAS coated glass slides (Matsunami Glass). Before staining, an antigen retrieval treatment was applied to the sections using 2100-Retiriever (Aptum Biologics) and R-BUFFER B (Electron Microscopy Sciences). The H&E staining protocol was as follows: (1) cell nuclei were stained by Mayer’s Hematoxylin Solution (FUJIFILM Wako Pure Chemical) for 5 min, (2) washed in water twice, (3) cytoplasm was stained by 0.5% Eosin Y Ethanol Solution (FUJIFILM Wako Pure Chemical) for 5 min, (4) washed in 100% ethanol three times, (5) sequentially treated in xylene for 5 min three times, (6) sealed by using Entellan New mounting medium (Merck).

All images (both bright field and fluorescence) were taken by THUNDER Imaging systems (Leica microsystems).

### Analysis of cell nuclei distribution for the tissues cultured using the 1-fiber HFB

DAPI staining images of the transverse tissue sections were first processed using ImageJ. In detail, the images were cropped using a 1 mm ×1 mm rectangular where the center of the rectangular matches with the center of the hollow fiber. Then, the hollow fiber is removed to avoid unwanted error counting of cell nuclei. Next, the images were thresholded and smoothed using the Guassian blur function. Next, analyzing particle functions were used to sort the cell nuclei and the coordinates of the nuclei were extracted and saved in .csv files. The .csv files were further processed using Python scripts to calculate the number of cell nuclei within the designated donut-shaped area.

### Analysis of myogenic marker-positive area under the differentiation condition

For the C2C12 tissues cultured using the 1-fiber HFB, α-actinin and myosin heavy chain expression images were first processed using ImageJ. In detail, the images were cropped using a 1 mm ×1 mm rectangular where the center of the rectangular matches with the center of the hollow fiber. Then, the hollow fiber is removed, and the images are thresholded. Next, the adjacent expression areas were merged and enclosed into a donut-shaped area, and the effective thickness of the donuts for both markers (MyHC and α-actinin) are extracted, calculated, and plotted using Python scripts.

For the chicken tissue cultured using the 50-fiber HFB, phase contrast images were first processed using ImageJ. In detail, each perfusable hollow was enclosed manually using the selection function. The hollows were then labeled according to different columns and rows. Each center position of the hollows was extracted and the distances between the adjacent hollows were analyzed using Python scripts. Next, cell nuclei and myosin heavy chain expression areas were enclosed with their coordinate and area extracted using ImageJ. Nuclei distribution and myosin heavy chain expression area were analyzed using Python scripts.

### Analysis of myotube orientation

For the chicken tissue cultured using the 50-fiber HFB, longitudinal sections staining images (α-actinin) were cropped around each hollow fiber to reduce the amount of calculation. Next, the hollow fibers and the myotubes surrounding the hollow fibers were enclosed and fitted using eclipses in ImageJ. After extracting the long-axis direction of all the fitted areas, the directional angles between the hollow fibers and their surrounding myotubes were calculated, analyzed and plotted using Python scripts.

### Analysis of muscle contraction of the tissues

Electric pulses (pulse duration: 20 ms, positive peak-to-peak period: 1 s for twitch contraction and 0.1 s for tetanic contraction, 18 Vpp) generated by a function generator, WF1968 (NF Corporation) were applied to the chicken muscle tissues cultured with perfusion (40 µL/min per hollow fiber, internal pressure: 38.1 mBar) or without perfusion (as a control) in the 50-fiber HFB for 9 days (N=7). The pulses were applied by placing a pair of gold electrodes at the diagonal corners (distance: ca. 1.73 cm) of the tissues. Since the cultured tissues have the tendency to shrink without the anchoring structure and will show deteriorated contractility, the tissues were stimulated on the HFB with the PDMS holder to keep the tension. The video was recorded by using an IX71 inverted microscope equipped with a DP74 camera, and imaging software cellSens (Evident). Using ImageJ software, the videos were cropped with its horizontal axis aligned to the length direction of the tissue, area of the tissue were extracted, and the effective width of the tissue were calculated for each frame.

### Texture analysis

The chicken skeletal muscle tissues cultured in 50-fiber HFB with (40 µL/min per hollow fiber) or without (as a control) perfusion for 9 days were used for texture analysis (N=1). Before the analysis, the tissue constructed by the perfusion culture was confirmed to have formed a contractile muscle tissue by applying electrical stimulation. After taking out from HFB and removing all hollow fibers, each tissue was cut into three equal parts (about 10 mm ×5 mm ×5 mm pieces). The force (g) to compress those raw tissue pieces to 50% thickness perpendicular to myofibers was measured by a compression test using texture analyzer, TA.XTplus (Eko Instruments) (measurement speed: 5 mm/sec, probe: φ35 mm cylinder).

### Amino acid analysis

Amino acid composition analysis by high-performance liquid chromatography (HPLC) was performed by Shimadzu corporation Analytical & Measuring Instruments Division (Kyoto, Japan). Briefly, the samples for HPLC were extracted with trichloroacetic acid solution from freeze-dried chicken muscle tissues cultured with (40 µL/min per hollow fiber) or without (as a control) perfusion in 50-fiber HFB for 9 days (N=2). Amino acids were separated using cation exchange column, Shim-pack Amino-Li (100 mm L. ×6.0 mm I.D) (Shimadzu) and detected by the post-column derivatization method using o-phthalaldehyde. For quantitative analysis, a calibration curve was created from the area value of amino acids mixture standard solution type H (FUJIFILM Wako Pure Chemical) added with Asparagine, Glutamine, and Tryptophan and diluted by lithium citrate buffer (pH 2.2). Each sample was analyzed three times and the mean values for two tissues under the same culture conditions were shown.

### Statistical analysis

We evaluated statistically significant differences by independent Student’s t-test or Tukey-Kramer test. All statistical analyses were performed using SciPy and Python statsmodels package. All data are represented as the average±SD.

## Acknowledgements

This work is partially supported by the JST-Mirai Program (Grant Number JPMJMI20C1), Japan. The authors would also like to thank Ms. Yuko Matsumura for technical assistance on animal experiments and HFB fabrication, Daicel Corporation for the courtesy of providing the hollow fibers, SAMCO Inc. for technical support on the water vapor-based plasma equipment, and Shimadzu corporation Analytical & Measuring Instruments Division for the courtesy of performing amino acid composition analysis of the cultured tissue samples.

## Author contributions

Conceptualization: M.N., A.S., and S.T.; methodology: M.N. and A.S.; investigation: M.N., A.S., and S.T.; visualization: M.N. and A.S.; funding acquisition: S.T.; project administration: S.T.; supervision: S.T.; writing – original draft: M.N. and A.S.; writing – review & editing: M.N., A.S., and S.T.

