## Supplementary material for "Centimeter-scale perfusable cultured meat with densely packed, highly aligned muscle fibers via hollow fiber bioreactor": SI.pdf

**Table S1. Calculation results for the internal pressure difference between the middle point of the hollow fiber and the culture medium outside ($P_m$)**

| HFB type | Flowrate (per fiber) | Hollow fiber half length | Outlet tube dia/length | Pressure drop on the outlet tube | Pressure drop on the hollow fiber (half length) | $P_m$ |
|---|---|---|---|---|---|---|
| **1-fiber** | 15 µL/min | 5 mm | 0.5 mm /15 mm | 0.02 | 0.09 | 0.1 mBar |
| **1-fiber** | 100 µL/min | 5 mm | 0.5 mm /15 mm | 0.12 | 0.63 | 0.8 mBar |
| **1-fiber** | 500 µL/min | 5 mm | 0.5 mm /15 mm | 0.59 | 3.13 | 3.7 mBar |
| **4-fiber** | 500 µL/min | 20 mm | 0.25 mm /15 mm | 37.55 | 12.52 | 50.1 mBar |
| **50-fiber** | 40 µL/min | 10 mm | 0.25 mm /15 mm | 37.55 | 0.5 | 38.1 mBar |

***Common parameters for the calculation:**

**Hollow fiber inner diameter:** 250 µm

**Viscosity:** 0.00072 Pa·s (medium with 10% FBS)

**Density:** 1000 kg/$m^3$

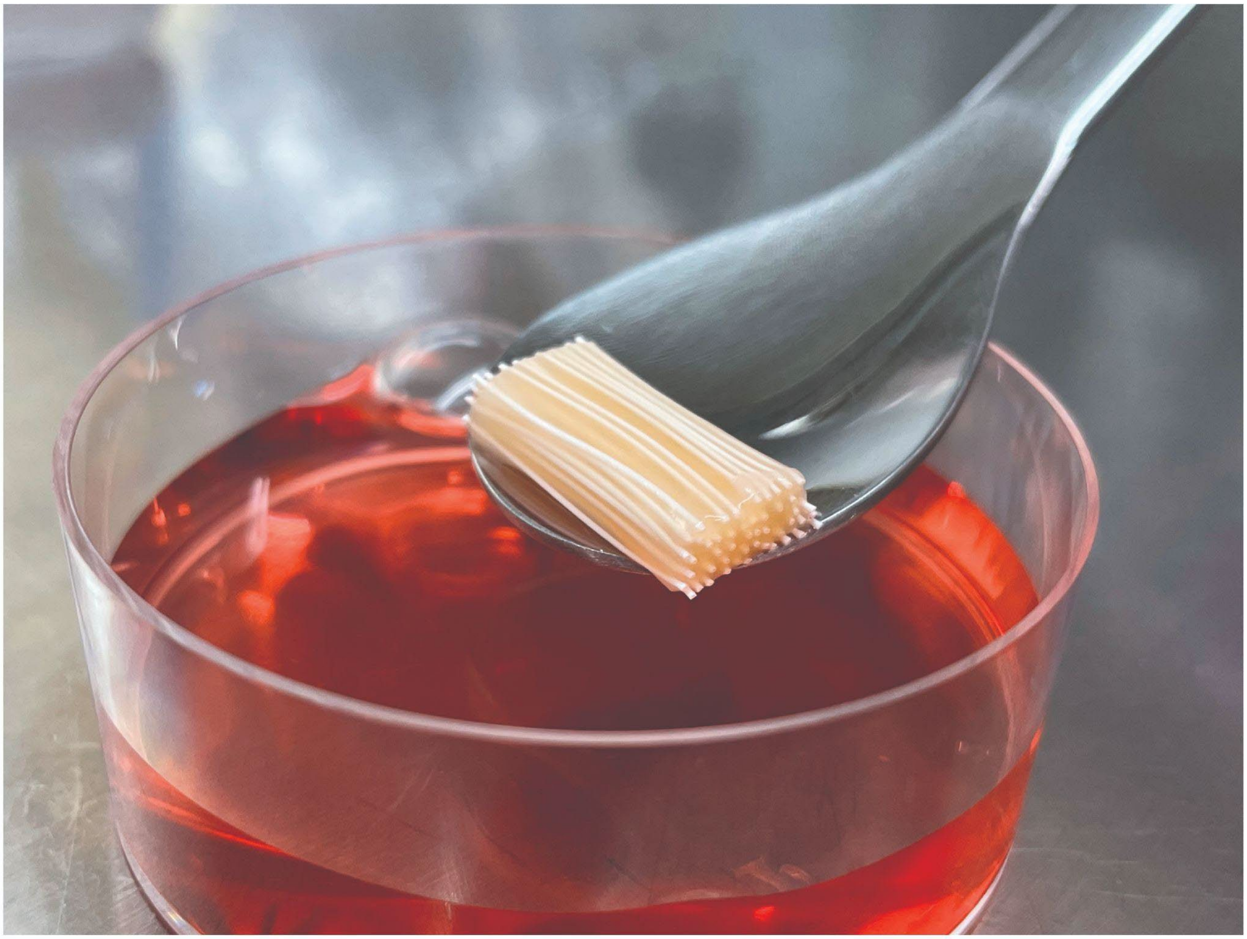

**Figure S1. Cultured chicken skeletal muscle tissue with embedded hollow fibers.**

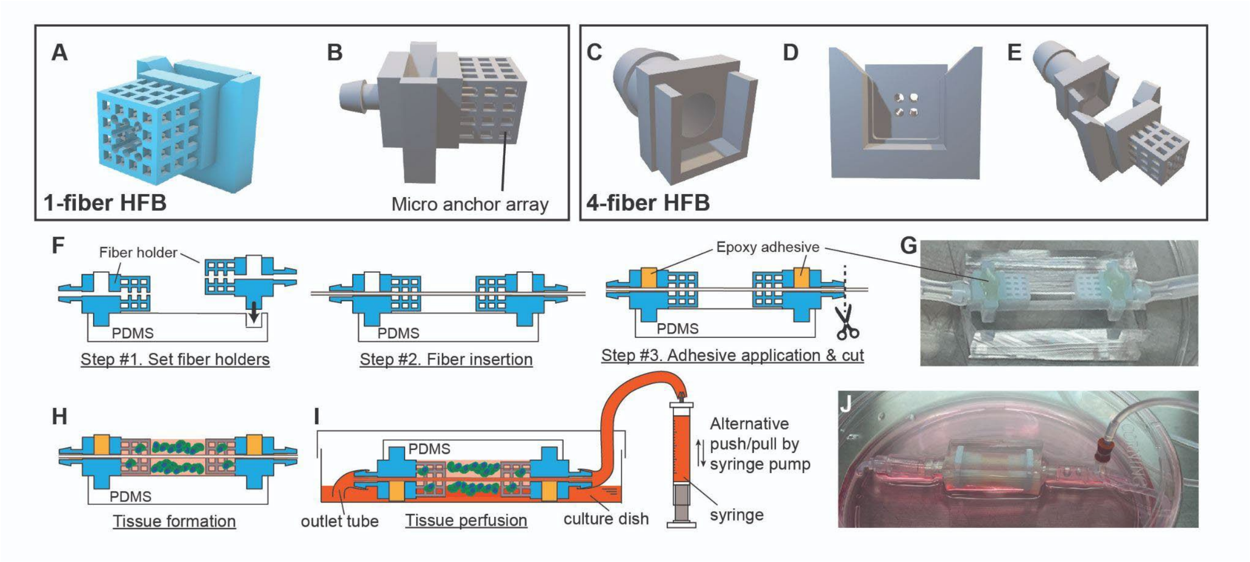


**Figure S2. Design, fabrication of the 1-fiber HFB, 4-fiber HFB and tissue culture setup. A-B.** *The design of the fiber holder for the 1-fiber HFB.* **C-E.** *The design of the fiber holder for the 4-fiber HFB. The fiber holder consists of two separate parts (***C** *and* **D***) and is assembled using glues after threading the hollow fibers.* **F.** *Detailed steps for the fabrication of the HFB (using 1-fiber HFB as an example).* **G.** *Photo of the fabricated 1-fiber HFB.* **H.** *Tissue formed by pouring hydrogel cell-suspension into the chamber of the HFB.* **I.** *After tissue formation, the HFB is flipped upside down, placed in a culture dish, and connected to a syringe through a via drilled on the lid of the dish. Alternative bidirectional flows are generated by the pull-push motion of the syringe pumps. The medium level of the dish is kept below 1 mm to ensure oxygen saturation level during the perfusion culture. The outlet tube of the HFB is dipped towards the bottom of the culture dish to ensure successful suction of culture medium during the pull phase of the syringe pump.* **J.** *Photo of the tissue (cultured using the 1-fiber HFB) during the perfusion culture.*

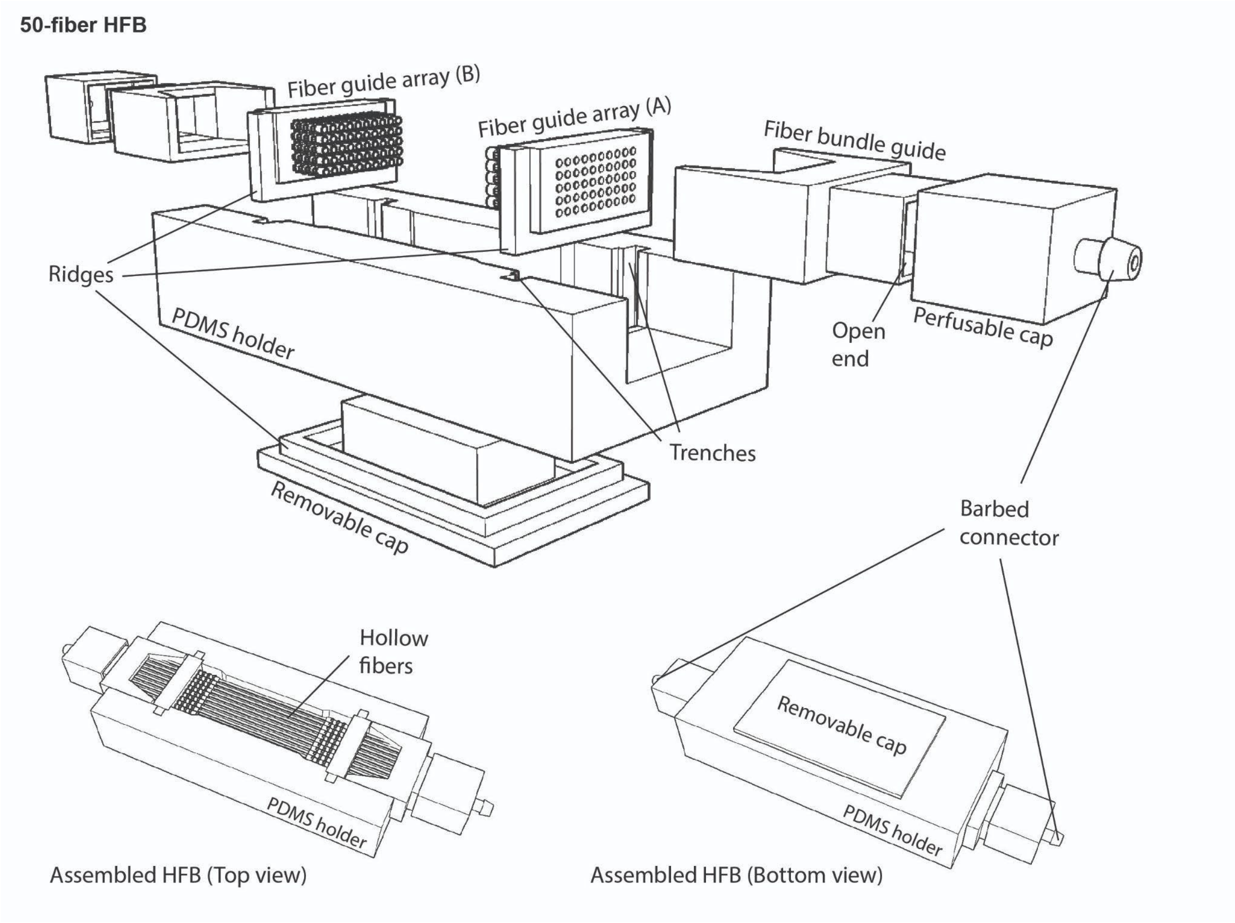


**Figure S3. Design, fabrication of the 50-fiber HFB.**

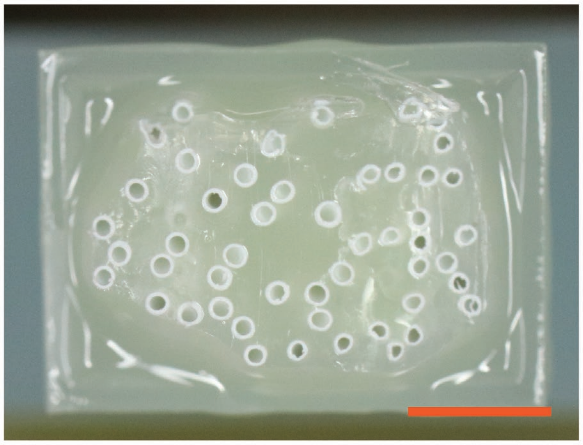

**Figure S4. Cross section of the cut hollow fibers (embedded in the adhesives) at the end of the fiber bundle guide. Scale bar, 2 mm.**
